# Deep residual neural networks resolve quartet molecular phylogenies

**DOI:** 10.1101/787168

**Authors:** Zhengting Zou, Hongjiu Zhang, Yuanfang Guan, Jianzhi Zhang

## Abstract

Phylogenetic inference is of fundamental importance to evolutionary as well as other fields of biology, and molecular sequences have emerged as the primary data for this task. Although many phylogenetic methods have been developed to explicitly take into account substitution models of sequence evolution, such methods could fail due to model misspecification and insufficiency, especially in the face of heterogeneities in substitution processes across sites and among lineages. In this study, we propose to infer topologies of four-taxon trees using deep residual neural networks, a machine learning approach needing no explicit modeling of the subject system and having a record of success in solving complex non-linear inference problems. We train residual networks on simulated protein sequence data with extensive amino acid substitution heterogeneities. We show that the well-trained residual network predictors can outperform existing state-of-the-art inference methods such as the maximum likelihood method on diverse simulated test data, especially under extensive substitution heterogeneities. Reassuringly, residual network predictors generally agree with existing methods in the trees inferred from real phylogenetic data with known or widely believed topologies. We conclude that deep learning represents a powerful new approach to phylogenetic reconstruction, especially when sequences evolve via heterogeneous substitution processes. We present our best trained predictor in a freely available program named *Phy*logenetics by *D*eep *L*earning (PhyDL, https://gitlab.com/ztzou/phydl).

## INTRODUCTION

A phylogeny is a tree structure depicting the evolutionary relationships among taxonomic units such as species, populations, or individuals (Nei and Kumar 2000; Felsenstein 2004; Yang 2006). Fully resolved phylogenies are typically binary, with pairs of taxa or groups of taxa joined hierarchically backward in time, representing historical splits and divergences between lineages. The central role of phylogenies in evolutionary biology is reflected in Charles Darwin’s Origin of Species, where the sole figure is a hypothetical phylogeny of some species (Darwin 1859). Apart from providing fundamental knowledge and framework for answering evolutionary questions (Romiguier, et al. 2013; Jarvis, et al. 2014; Lamichhaney, et al. 2015; Simion, et al. 2017), the concept of phylogeny is widely used in other fields such as developmental biology (Kalinka, et al. 2010), cell lineage reconstruction (Salipante and Horwitz 2006), epidemiology (Cassan, et al. 2016), cancer biology (Cooper, et al. 2015; Leung, et al. 2017), wildlife conservation (Carvalho, et al. 2017), forensics (Metzker, et al. 2002; Bhattacharya 2014), and even linguistics (Dunn, et al. 2005; Atkinson, et al. 2008).

Despite their broad importance, the underlying phylogenies of existing species or taxonomic units (taxa) are not directly visible, and therefore need to be inferred. With the rapid accumulation of sequenced DNAs over the last few decades, alignments of DNA or protein sequences from extant species have emerged as the primary data for phylogenetic inference. Over the past 50 years, multiple tree inference methods have been developed, including for example, Unweighted Pair Group Method with Arithmetic Mean (UPGMA), Minimum Evolution (ME), Neighbor-Joining (NJ), Maximum Parsimony (MP), Maximum Likelihood (ML), and Bayesian Inference (BI). Among them, distance-based methods such as UPGMA, ME, and NJ first estimate evolutionary distances for all pairs of taxa, and then infer the best tree under various criteria about branch lengths; character-based methods directly fit the observed sequence data to each tree in the topological space, and search for the best tree that can explain the data by the smallest number of substitutions (MP), the highest likelihood (ML), or the highest posterior probability (BI) (Nei and Kumar 2000; Felsenstein 2004; Yang 2006).

In the computation of evolutionary distances, likelihood values, or probabilities, explicit Markovian models of nucleotide or amino acid substitution are usually adopted for each nucleotide or amino acid site. The transition rate matrix ***Q*** describes the rate with which one state changes to another, and is composed of a symmetric exchangeability matrix ***S*** defining the tendency of exchange between each pair of states and a vector ***π*** denoting equilibrium frequencies of all states. In practice, researchers almost universally assume that different sites in a sequence evolve independently but follow the same ***Q*** matrix or one of a few ***Q*** matrices. Evolutionary rate variation among sites is typically modeled by multiplying tree branch length *t* with a gamma-distributed factor *r* (Nei and Kumar 2000; Felsenstein 2004; Yang 2006). This is undoubtedly a gross oversimplification of actual evolution, because different sites in a gene or protein play different structural and functional roles and are subject to different constraints and selections. In addition, the site-specific constraints and selections (e.g., ***π*** and *r*) can vary with time in the course of evolution (Fitch and Markowitz 1970; Mooers and Holmes 2000; Penny, et al. 2001; Tarrío, et al. 2001; Lopez, et al. 2002; Zou and Zhang 2015). Hence, there are both site and time heterogeneities in sequence evolution.

Theoretically, the ME and ML methods are guaranteed to yield the correct tree when (i) the model of sequence evolution used is correct and (ii) the sequences are infinitely long (Nei and Kumar 2000; Felsenstein 2004; Yang 2006). While the second criterion can be more or less satisfied by today’s large genomic data, the first cannot because the model of sequence evolution is almost always unknown. Even though it is in principle possible to estimate evolutionary models from an alignment, the estimation depends on having a correct phylogeny, which is unknown. Making the situation worse is the fact that an increase in data size makes phylogenetic inference more sensitive to model assumptions, because a slight model misspecification can lead to erroneous results with high statistical support (Lemmon, et al. 2013).

Although there is a continuing effort to make models more realistic (Ronquist and Huelsenbeck 2003; Stamatakis 2014), model misspecification is and will likely be the norm rather than the exception. Computer simulations have repeatedly shown that model misspecification can cause gross errors in phylogenetic inference (Takezaki and Gojobori 1999; Nei and Kumar 2000; Felsenstein 2004). Specifically, different treatments of site and time heterogeneities in sequence evolution affect tree inference in many case studies (Lockhart, et al. 1992; Foster and Hickey 1999; Tarrío, et al. 2001; Roure and Philippe 2011; Feuda, et al. 2017). While there have been efforts to incorporate such heterogeneities into phylogenetics, highly heterogeneous evolutionary processes, especially time heterogeneity, are difficult to model (Lake 1994; Foster 2004; Lartillot, et al. 2009; Heaps, et al. 2014).

Given the above difficulty, we propose to use deep neural networks, a machine learning approach, for phylogenetic inference, because this general-purpose approach requires no explicit modeling of the subject process of inference and enables learning properties and patterns of interest from data without knowing them *a priori* (Franklin 2005; Murphy 2012). A deep neural network predictor consists of a cascade of linear functions (vector product) and nonlinear activation functions. Given training data with known answers, a training process can perform gradient descent optimization on the coefficients such that the outcome of the updated network approximates the answer. The method has been successfully applied to a wide range of inference tasks including image classification, speech recognition, natural language processing, and functional annotations of genome sequences (Graves, et al. 2013; Luong, et al. 2015; Szegedy, et al. 2015; Zhou and Troyanskaya 2015; He, et al. 2016).

A commonly used class of deep neural network is the convolutional neural network (CNN). The core layers in a CNN perform convolution or dot-product of a coefficient matrix and one patch of data repeatedly to the entire input, resulting in a matrix to be used for further convolution in the next layer or the output of the whole network. Recent successes in applying CNN to genome sequence analysis has demonstrated its great potential in identifying sequence patterns (Zhou and Troyanskaya 2015). In this study, we formulate a residual neural network, a type of CNN with proven success in image classification, to predict the topology of trees with four taxa (i.e., quartet trees). We train this model on huge datasets generated by simulating protein sequence evolution with extensive site and time heterogeneities. The resulting residual network predictors were evaluated against current phylogenetic inference methods on both simulated test datasets and real phylogenetic datasets. We show that our predictors are generally superior to the existing methods, especially on difficult quartet trees under heterogeneous evolutionary schemes. We present our predictor as *Ph*ylogenetics by *D*eep *L*earning (PhyDL, https://gitlab.com/ztzou/phydl).

## RESULTS

### Construction and training of residual network predictors for phylogenetics

Our deep neural network predictor resembles the original residual network proposed for image classification (He, et al. 2016) but uses one-dimension (1-D) convolution instead of 2-D (**Table S1**, see Materials and Methods). The predictor takes an alignment of four one-hot-encoded amino acid sequences with an arbitrary length as input, which contains 4 (taxa) × 20 (amino acids) channels. The predictor applies two rounds of convolution which reduces the number of input channels to 32, followed by a series of residual modules and pooling layers. Each residual module consists of two sequential rounds of 1-D convolution + 1-D batch-normalization + rectified linear unit (ReLU) activation, results of which are added back to the input of the residual module as its output. Following four groups of residual modules and pooling layers, a dense layer of linear functions transforms the data to three probabilities, respectively representing the likelihood of each of the three possible tree topologies regarding the four taxa concerned. The fact that the residual module “remembers” the input data by adding it directly to convolutional output is expected to guard against the “vanishing gradients” effect in very deep networks (He, et al. 2016).

Few cases exist where we know the true phylogeny of taxa with sequence data, but deep neural networks need large amount of truth-labeled data for training. Hence, we simulated Markovian amino acid sequence evolution along randomly generated trees as training data (see Materials and Methods). If all possible values of parameters of a quartet tree and associated sequence data form a parameter space, our sampling should be diverse and heterogeneous enough to contain and represent the subspace of real phylogenetic data. We generated random trees with more than four taxa and simulated amino acid sequences (of varying lengths) according to the trees. The amino acid exchangeability matrix ***S*** of each simulated alignment is randomly chosen from nine widely used empirical matrices specified in PAML 4.9 (Yang 2007). Regarding site heterogeneity, a relative evolutionary rate *r* was assigned to each site, along with a site-specific amino acid equilibrium frequency profile ***π*** drawn from large alignments of real nuclear and organelle genome-encoded proteins. Each branch in the tree has a certain probability of containing time heterogeneity, which is realized by shuffling the *r* values among some sites (rate swap) and changing the equilibrium profile ***π*** of each site (see Materials and Methods). After the generation of each tree and associated sequence data, we pruned the tree so that only four taxa remain, hence creating a quartet tree sample ready for training, validation, or testing of the residual network predictor. In quartet tree inference, a traditional challenge involves trees with two long branches, each grouped with a short branch and separated by a short internal branch. This type of trees is subject to long-branch attraction (LBA), which erroneously clusters the two long branches as a result of random sequence convergence (Felsenstein 1978). LBA is especially problematic for MP, but also occurs to other methods under misspecified evolutionary models. To ensure that the training process involved diverse and challenging learning materials, we pruned a proportion of trees to four randomly chosen taxa (normal trees), and the other trees to four taxa with high LBA susceptibility (LBA trees, see Materials and Methods for detailed criteria).

Training consisted of multiple iterative epochs, based on a total training pool of 100,000 quartets containing 85% normal trees and 15% LBA trees. Cross-entropy loss was used to measure the disagreement between residual network predictions and true tree topologies (see Materials and Methods for the mathematical definition). In each epoch, 2,000 training samples were randomly drawn from the total pool of training data; the residual network predictor updated itself according to backpropagation from cross-entropy loss on the training samples (training loss), resulting in a new predictor. After each epoch, performance of the current predictor was evaluated by its cross-entropy loss (validation loss) in predicting 2,000 validation samples, which were separately simulated but under the identical schemes as the training data. After 500 epochs, the training process would continue except when there had been 80 epochs with higher validation loss than the current minimum loss among all epochs. To investigate the impact of training data on the performance of the resultant predictor, we trained our residual network on three datasets generated under different simulation schemes, and consequently obtained three series of predictors named deep neural network 1 (dnn1), dnn2, and dnn3. The training data used for dnn1 (training1 in **Table S2**) contained more time heterogeneities than those for dnn2 (training2 in **Table S2**) and dnn3 (training3 in **Table S2**). Additionally, training trees for dnn2 have gamma-distributed branch lengths shorter than those uniformly distributed ones for dnn1 and dnn3 on average (**Table S2**; see Materials and Methods). Training of dnn1, dnn2, and dnn3 stopped at 669, 1359, 1179 epochs, respectively, all reaching low levels of validation loss around 0.15 to 0.3 (**Fig. 1a**).

**Fig. 1.**
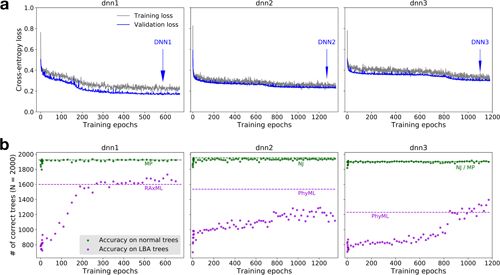
Residual networks gain predictive power in resolving quartet trees through training. (**a**) Cross-entropy loss of the predictors in training and validation after each training epoch. Blue arrows indicate the predictors used in subsequent analyses, because these predictors have the lowest validation cross-entropy losses. (**b**) Performances of residual networks at sampled epochs on test datasets with normal trees and LBA trees, respectively. Dashed lines indicate the best performances among existing methods examined on normal trees or LBA tress, respectively.

### Performance of the residual network predictors on test data

We first examined the performance of the residual network predictors on sequence data that were simulated using the same parameters as in the generation of the corresponding training data. Because the training data were a mixture of normal and LBA trees, we evaluated prediction accuracies of dnn1, dnn2, and dnn3 at different training time points, on separate test datasets with only normal trees (testing1_nolba, testing2_nolba, and testing3_nolba in **Table S2**, green dots in **Fig. 1b**) and only LBA trees (testing1_lba, testing2_lba, and testing3_lba in **Table S2**, violet dots in **Fig. 1b**). Interestingly, the predictors gained optimal performance on normal trees within 10 epochs during training, but improved much more slowly in predicting LBA trees (**Fig. 1a**).

To benchmark the performances of the residual network predictors, we compared them with several widely used phylogenetic inference methods: NJ and MP implemented in the software MEGA X (Kumar, et al. 2018), ML implemented by the software RAxML v8 (Stamatakis 2014) and PhyML v3.1 (Guindon, et al. 2010), and BI implemented in MrBayes 3.2 (Ronquist, et al. 2012). We used reasonable parameter settings in each of these programs to ensure fair comparisons (see Materials and Methods). For example, pairwise distances in NJ were calculated with the Jones-Taylor-Thornton (JTT) model and gamma-distributed rate variation (shape parameter = 1). In MP, the subtree-pruning-regrafting (SPR) branch-swapping algorithm was used in searching for the optimal tree. In RAxML, the model “PROTCATAUTO” was used so that the program infers trees under different substitution matrices and reports the best result. The LG substitution matrix (Le and Gascuel 2008) was specified in both PhyML and MrBayes. In MrBayes, two replicate runs of four 20,000-step MCMC sampling were performed for each sequence alignment, and the consensus trees were summarized after the 25% burn-in steps (see Materials and Methods for details).

At later stages of training, both dnn1 and dnn3 showed performance closely matching or superior to the best current inference methods on normal and LBA trees (dashed lines in **Fig. 1b**). The predictor dnn2 showed poorer performance than the best performance of existing methods, especially when compared with PhyML on LBA trees, probably due to the fact that it was trained mainly on trees with relatively short branches. The three predictors at epochs with the lowest validation loss (epoch 588 for dnn1, 1272 for dnn2, and 1098 for dnn3), referred to as DNN1, DNN2, and DNN3, respectively, were used in subsequently analyses.

A detailed comparison showed that, although the accuracies of all methods examined were quite high, the residual network predictors generally outperformed the traditional methods (**Table 1**). For example, of the 2,000 test samples (testing1_mixed) simulated and mixed under the same parameters as the DNN1 training data (training1), DNN1 correctly predicted 1,881 trees, whereas the best performance of any existing method made 1868 correct predictions (RAxML). On the normal trees (testing1_nolba), both DNN1 (1,926 correct predictions) and DNN3 (1,935 correct predictions) surpassed all existing methods examined, among which the best performance was 1,924 correct trees by MP. For LBA trees (testing1_lba), DNN1 inferred 1,659 correct trees, while the best performance of any existing method was only 1,600 correct trees (RAxML). The same trend was observed on test data simulated using parameters used in the generation of the training data of DNN2 and DNN3, respectively. Overall, both DNN1 and DNN3 showed superior performance in at least six of the nine test datasets considered when compared with any existing method examined here (**Table 1**). Interestingly, DNN2 did not perform well even on test datasets similarly simulated as its training data (**Table 1**), consistent with the earlier observation (**Fig. 1b**).

**Table 1.** Numbers of correctly inferred quartet trees by residual network predictors and existing methods on test datasets simulated under the training simulation schemes.

| Test datasets | # of test trees | DNN1 | DNN2 | DNN3 | NJ | MP | RAxML | PhyML | MrBayes |
| --- | --- | --- | --- | --- | --- | --- | --- | --- | --- |
| testing1_mixed<sup>a</sup> | 2000 | **1881**<sup>b</sup> | 1810 | 1851 | 1844 | 1791 | 1868 | 1860 | 1860 |
| testing1_nolba | 2000 | 1926 | 1910 | **1935** | 1910 | 1924 | 1912 | 1896 | 1906 |
| testing1_lba | 2000 | **1659** | 1152 | 1446 | 1416 | 1078 | 1600 | 1592 | 1475 |
| testing2_mixed<sup>a</sup> | 2000 | **1882** | 1831 | 1860 | 1868 | 1807 | 1853 | 1841 | 1842 |
| testing2_nolba | 2000 | 1941 | 1931 | 1948 | **1951** | 1933 | 1926 | 1917 | 1920 |
| testing2_lba | 2000 | **1611** | 1254 | 1512 | 1437 | 1045 | 1494 | 1536 | 1479 |
| testing3_mixed<sup>a</sup> | 2000 | 1792 | 1733 | **1794** | 1753 | 1736 | 1758 | 1731 | 1738 |
| testing3_nolba | 2000 | 1900 | 1897 | **1913** | 1904 | 1904 | 1890 | 1867 | 1879 |
| testing3_lba | 2000 | **1306** | 986 | 1286 | 1140 | 867 | 1190 | 1230 | 1162 |

<sup>a</sup> “\_mixed” indicates the test set contains 85% normal trees and 15% LBA trees (same as in the training data).
<sup>b</sup> The bold cell in each row indicates the best performance among all methods.

### Performance of the residual network predictors on diverse simulated data

Although our training data are heterogeneous, they represent but a part of the tree parameter space. As a result, our predictors may perform well only on test data similarly generated as the training data. To examine this possibility, we investigated the performance of our predictors when certain tree properties vary greatly. Three series of test datasets were simulated with varying tree depths, sequence lengths, and heterogeneity levels.

In the first series of six datasets, one thousand 20-taxon trees were simulated with individual branch lengths in the range of [0.02, 0.2), [0.2, 0.4), [0.4, 0.6), [0.6, 0.8), [0.8, 1.0), and [1.0, 2.0), respectively, before being pruned to quartet samples; all other parameters were unchanged (test_branch_00 – test_branch_05 in **Table S2**). As expected, all three residual network predictors and five existing methods show the best performances when branches are not too short nor too long (**Fig. 2a**). Furthermore, DNN1 and DNN3 are more accurate than all existing methods examined, regardless of the branch lengths (**Fig. 2a**). The performance of DNN2 is generally comparable to those of the best existing methods, although it becomes inferior when tree branches are long (in the range [1, 2)) (**Fig. 2a**), probably because DNN2 was trained mainly on trees with short branch lengths.

**Fig. 2.**
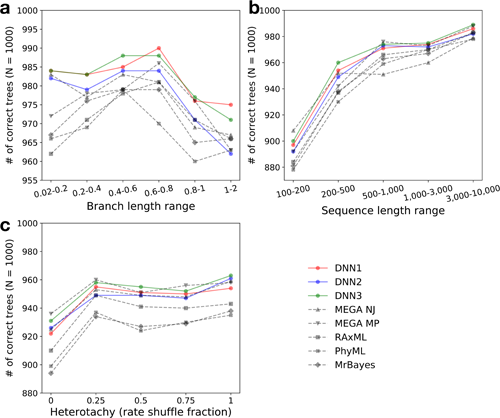
Residual network predictors generally outperform existing methods on quartet trees with diverse properties. Numbers of correct inferences out of 1,000 trees are shown for different test datasets with (**a**) different ranges of branch lengths, (**b**) different ranges of amino acid sequence lengths, and (**c**) different levels of heterotachy.

Second, we simulated five 1,000-tree datasets with the sequence length ranging from [100, 200) to [3,000, 10,000) amino acids (test_seqlen_00 – test_seqlen_04 in **Table S2**). As expected, the accuracy of every inference method increases with sequence length, from approximately 900 to 980 correct trees (**Fig. 2b**). No single existing method shows higher accuracies than DNN1 or DNN3 on more than two datasets, indicating that our residual network predictors are superior at various sequence lengths.

Third, we varied the level of heterogeneity in evolution. The evolutionary rate variation of the same site among different tree branches, or heterotachy, was realized by shuffling *r* values among a proportion (*p*) of amino acid sites at the beginning of each branch. We simulated five sets of 1,000 trees with *p* ranging from 0 to 1 (test_heterotachy_00 – test_heterotachy_04 in **Table S2**). The accuracy of DNN3 is comparable to the best performance of any existing method, especially when the heterotachy level is high (e.g., 963 correct trees by DNN3 versus 959 by NJ when *p* = 1), while DNN1 and DNN2 outperform all existing methods except NJ (**Fig. 2c**). Thus, our predictors work well under various degrees of heterotachy.

### Performance of the residual network predictors on LBA data with heterotachy

As mentioned, LBA trees represent a group of difficult cases in phylogenetic inference. Previous studies reported that different phylogenetic methods show different sensitivities to LBA when various levels of heterotachy exist (Kolaczkowski and Thornton 2004; Philippe, et al. 2005). Because our residual network predictors were trained on heterogeneous sequence data, they may be less sensitive than other methods to LBA in the presence of heterotachy. To this end, we simulated two series of quartet tree datasets with directly assigned branch lengths: the two short external branches have lengths *b* ranging from 0.1 to 1.0, the two long branches have lengths *a* ranging from 2*b* to 40*b*, and the internal branch has a length *c* ranging from 0.01*b* to 1*b* (**Fig. 3a**). Each dataset contains 100 simulated quartet trees. The first series of trees were simulated with no time heterogeneity (testlba_F_h0 in **Table S2**), while the second series were simulated with heterotachy (shuffling site rate *r*, testlba_F_h1 in **Table S2**). We then evaluated the performances of our residual network predictors and the existing methods on these two series of test datasets. Accuracies of all inference methods decrease when *b* increases (**Fig. 3b**), *a*/*b* ratio increases (**Fig. 3b**), or *c*/*b* ratio decreases (**Fig. 3c**). As expected, MP almost always exhibits the worst performance under LBA-susceptible conditions (**Fig. 3c**, **Fig. S1**). When *a*/*b* is large (e.g., last row in **Fig. 3c** and **Fig. S1**), almost all methods produce wrong trees, whereas the opposite is true when this ratio is small (e.g., first row in **Fig. 3c** and **Fig. S1**). Between these two extremes of tree parameter space, residual network predictors generally perform comparably or slightly inferior to PhyML and MyBayes when there is no heterotachy in evolution (**Fig. S1**). However, on the series of trees with heterotachy, DNN2 and DNN3 outperform the existing methods in many parameter combinations (indicated by colored pentagons in **Fig. 3c**). Because heterogeneity is prevalent in actual sequence evolution, these results suggest practical utilities of our predictors.

**Fig. 3.**
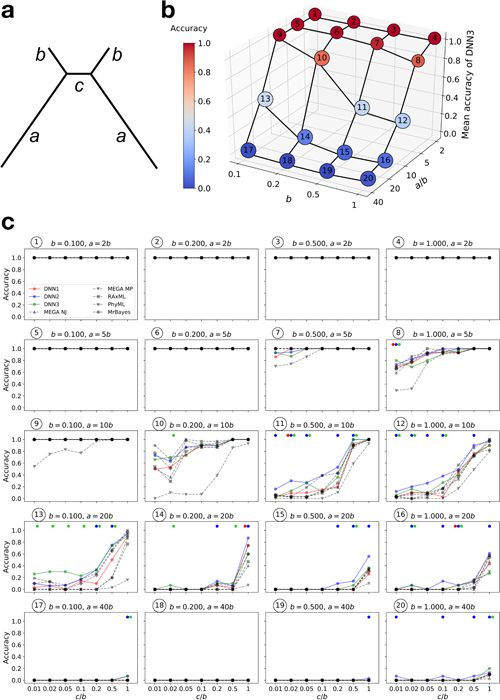
Residual network predictors generally outperform existing methods on LBA trees with heterotachy. (**a**) A schematic quartet tree showing branch length notations. (**b**) A 3-D surface of mean DNN3 accuracies (also indicated by color) across all *c*/*b* levels in each subplot of panel (c), in the space of *b* levels and *a*/*b* levels. Circled numbers correspond to those in (c). (**c**) Proportions of 100 quartet trees correctly inferred by our predictors (shown by different colors) and the existing methods (shown by different grey symbols) under various combinations of the parameters *b, a*/*b*, and *c*/*b*. For each *c/b* level indicated on the *X*-axis, if a residual network predictor performs better than all existing methods, a pentagon of the corresponding color is drawn on the top of the panel.

### Residual network predictors generally support accepted topologies for actual data

Although the residual network predictors, especially DNN1 and DNN3, perform well on simulated sequence data, their performance on actual data is unknown. Two types of actual data exist. In the first type, the true tree is known; consequently, the performances of various tree building methods can be objectively compared. But this type of data is extremely rare and hence one cannot draw general conclusions on the basis of these data. In the second type, a widely believed tree exists even though it is not guaranteed to be the true tree. Because the widely believed tree could not possibly be widely believed if it were not strongly supported by some existing methods, residual network predictors cannot outperform these existing methods on such data. Even with these serious caveats, the real data allow a sanity check of our predictors that were trained exclusively on simulated data.

We first tested our predictors using an alignment of 19 red fluorescent protein (FP) sequences (with a length of 225 amino acids) that were generated through experimental evolution (via error-prone PCR) in the lab (Randall, et al. 2016). Hence, the true tree of the 19 sequences are known (**Fig. S2a**). Of 1,000 quartet trees randomly pruned from the 19-taxon tree, DNN3 correctly inferred 800 trees. NJ and MP respectively outperform DNN3 by 18 and 10 correct inferences, while PhyML and MyBayes have worse performances than all three residual network predictors (**Table 2**). Hence, our predictors show comparable performances as the existing methods. The short sequences and the artificial substitution process resulting from error-prone PCR may be partly responsible for the lower performance of ML and BI than NJ and MP.

**Table 2.**
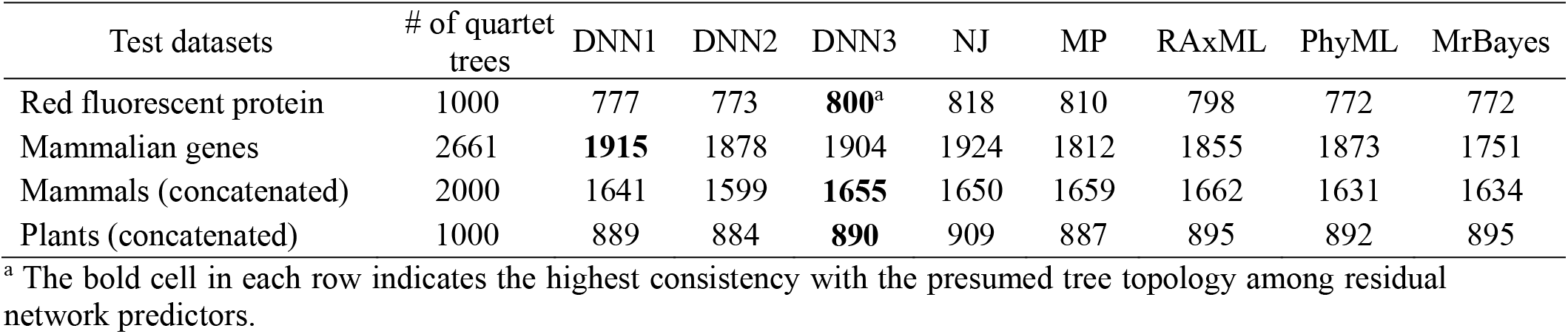
Numbers of inferred quartet trees by residual network predictors and existing methods that are consistent with the presumed tree topologies in actual phylogenetic data.

Next, we compiled test datasets based on the mammalian phylogeny. According to previous studies, we used a tree of 24 mammals as the presumed true tree (**Fig. S2b**), avoiding major controversial nodes such as the relationships among four eutherian superorders and those among the four orders within Laurasiatheria (Romiguier, et al. 2013). A total of 2,684 genes with filtered amino acid sequences for the 24 species were downloaded from OrthoMaM v10 (Scornavacca, et al. 2019). We conducted two different tests. In the first test, ungapped alignments with more than 50 amino acids for four randomly selected species were collected as a test dataset, totaling 2,661 genes. Most predictions of DNN1 (1915 consistent trees) and DNN3 (1904 consistent trees) are consistent with the original topology; they are slightly inferior to NJ (1924 consistent trees) but are superior to the other existing methods (“Mammalian genes” in **Table 2**). In the second test, we concatenated the sequences of all 2,684 genes and randomly sampled 2,000 four-species alignments of 100–3,000 amino acid sites. In this test, DNN1 (1641 consistent trees) and DNN3 (1655 consistent trees) show consistency levels within the range of existing methods (1631–1662 consistent trees). That is, they are less consistent with the presumed true tree than NJ, MP, and RAxML, but more consistent than PhyML and MrBayes (“Mammals (concatenated)” in **Table 2**).

Third, we examined the phylogeny of seed plants (spermatophytes). We compiled a highly multifurcating tree of 25 plant species (**Fig. S2c**), only retaining the bifurcating nodes of seven large taxonomic groups: gymnosperms, ANA grade, monocots, fabids, malvids, campanulids, and lamiids (Wickett, et al. 2014; Byng, et al. 2016). One thousand four-species alignments of 100–3,000 amino acid sites were randomly sampled from a concatenated alignment of 604 genes (Wickett, et al. 2014). For this test dataset, DNN1, DNN3, and all existing methods except NJ inferred 887–895 trees that are consistent with the presumed true tree; NJ inferred 909 consistent trees (**Table 2**).

With the above results from analyzing three real datasets, we conclude that the residual network predictors pass the sanity check.

## DISCUSSION

We have constructed the first deep residual neural networks for the task of inferring quartet phylogenies from amino acid sequence alignments. We trained these residual networks on simulated sequence data and showed that the trained predictors perform well on testing data similarly simulated. We found that our residual network predictors compare favorably with state-of-the-art phylogenetic methods under a variety of different sequence evolution schemes and tree parameter ranges. Specifically, our predictors outperform all examined existing methods on difficult LBA quartet trees involving extensive site and time heterogeneities in evolution. The sanity check using real phylogenetic datasets validates our predictors and reveals no sign of overfitting to the training schemes. Thus, training residual neural networks on heterogeneous phylogenetic data generated by computer simulation proves to be a promising solution to the current difficulty in this field. Based on the performances in all analyses, we have formulated DNN3 into a ready-to-use phylogenic inference program named *Ph*ylogenetics by *D*eep *L*earning (PhyDL, https://gitlab.com/ztzou/phydl).

The training processes provide interesting information on how residual networks gradually learn to extract phylogenetic signals from sequence alignments. In dnn1’s training process, while the ability to infer normal quartet trees was quickly gained in 10 epochs, the ability to resolve LBA trees was gained much more slowly (**Fig. 1a**). For instance, the ability of dnn1 to resolve LBA trees was no greater than that of the MP method (1078 correct trees per 2,000 trees tested) before 100 epochs. The conspicuous improvement in dnn1’s LBA-resolving ability occurred between 100 and 200 epochs (**Fig. 1b**). In dnn3’s training process, the accuracy in resolving LBA trees even stayed around the level of MP (867 correct trees per 2,000 trees tested) for approximately 700 epochs before its rapid increase and eventual surpass of the highest level among all existing methods, 1230 correct trees out of 2,000 tests by PhyML (**Fig. 1b**). These step-wise learning patterns suggest the possibility that, residual networks first captured straightforward phylogenetic signals that can also be picked up by simple methods such as MP, and then learned to extract signals of complex Markovian evolutionary processes.

Despite the simple optimizing criteria of MP and NJ, for most normal tree datasets of this study, these two methods outperform ML and BI even when the evolutionary process is relatively complex. This is probably because alignments with four taxa do not provide sufficient information for accurate estimation of parameters of complex evolutionary models, which could occur in reality. In this sense, our residual network predictors are advantageous in that they do not rely on the focal data to parameterize the model. Furthermore, commonly used ML and BI programs do not consider time heterogeneity in sequence evolution; consequently, their performance may be severely impaired due to model misspecification or insufficiency in the face of extensive heterogeneities. By contrast, deep learning allows a predictor to acquire the ability to infer trees even when the sequence evolution is highly heterogeneous. More importantly, residual network predictors require no *a priori* specification of mechanistic sequence evolution models during training, relieving the risk of model misspecification.

Apart from its inference accuracy, deep neural networks running on graphics processing units (GPUs) are also time-efficient when inferring trees. As a prediction speed benchmark, we inferred the topology of 100 simulated quartet trees with sequence length of 2,000 amino acids and all branch lengths equal to 1 (testtime_01 in **Table S2**). On an Intel Xeon W-2133 central processing unit (CPU) core (3.6 GHz), the NJ and MP algorithms implemented in MEGA spent 0.077s and 0.127s, respectively, for an average tree, while RAxML, PhyML, and MrBayes spent 2.81s, 7.82s, and 41.7s, respectively. On the same CPU core, residual network predictors on average use 1.09–1.20s per tree, slower than NJ and MP but faster than ML and BI. However, using the CPU core alongside a Nvidia Titan Xp graphical card, our three predictors spent 0.054–0.055s per tree, faster than any existing method compared here. The total time cost for 100 trees ranged from 9.7s to 10.0s for our predictors, including the hang-over time of loading the predictors onto GPU. The utilization of GPU-accelerated calculation by deep neural networks can achieve high efficiency in massive phylogenetic inference tasks, either on public servers or on properly configured personal computers.

Despite the generally high accuracy, our residual network predictors fail to surpass the performance of the best existing methods under some conditions. However, in practice, it is unknown which existing method performs the best for a given dataset because the underlying true tree is unknown. In our analyses, for instance, NJ and MP achieve good performance on normal trees, but show lower accuracies on LBA trees, while the opposite is true for ML. That our residual network predictors have an overall superior performance suggests that it can be used for diverse quartet tree inference tasks.

Recent years have seen multiple deep learning applications in population and evolutionary genetics (Sheehan and Song 2016; Kern and Schrider 2018). When we were preparing this manuscript, Suvorov and colleagues reported the implementation and training of a deep convolutional neural network for inferring quartet trees from DNA sequences (Suvorov, et al. 2019). Because our predictor was trained on protein sequences while theirs was trained on DNA sequences, the performances of the two predictors cannot be directly compared. Nevertheless, several differences are apparent. First, we used residual neural networks, which, compared with the traditional convolutional neural network used by Suvorov et al, allow deeper network structures without suffering from the “vanishing gradients” effect, hence can potentially achieve better learning of complex evolution processes (He, et al. 2016). In fact, our networks have 16 layers of convolution while Suvorov et al.’s networks have eight. Second, Suvorov et al. simulated gapped and ungapped alignments and showed that the advantage of their predictor over existing methods is mainly in dealing with gaps. We simulated only ungapped alignments and found that our predictor outperforms existing methods even in the absence of gaps. Third and most importantly, our predictor performs well not only on simulated but also on real, diverse phylogenetic data, while Suvorov et al.’s predictor has yet to be evaluated on real data. This difference is especially relevant because real data have site and time heterogeneities as well as sequence length variations, which were included in our but not Suvorov et al.’s training data. Note that the implementation of residual neural networks for resolving quartet trees is readily applicable to nucleotide sequence data, and it will be of interest to develop residual network predictors for nucleotide sequences in the near future.

Quartet trees are the smallest possible trees, and here we treated quartet tree inference as a classification task because only three possible tree topologies exist for four taxa. However, with an arbitrary number of *N* taxa, the possible number of topologies can be astronomical. Hence, a structure prediction methodology must be applied, which requires more complicated network formulations than convolutional neural networks (Joachims, et al. 2009). Given the success of deep learning for quartet tree inference demonstrated here, we are cautiously optimistic that machine learning will improve phylogenetic reconstruction in general and thus call for more studies in this promising area.

## MATERIALS AND METHODS

### Residual network structure and training

The residual neural network is implemented using the python package PyTorch v1.0.0. The raw input data, as in conventional phylogenetic inference software, are four aligned amino acid sequences (denoted as taxon0, taxon1, taxon2, and taxon3, hence dimension 4×*L*). This is then one-hot-encoded, expanding each amino acid position into a dimension with 20 0/1 codes indicating which amino acid is in this position. The 4×20×*L* tensor is transformed to an 80×*L* matrix and fed into the residual network described in **Table S1** with eight residual modules. The output of the network includes three numbers representing the likelihood that taxon0 is a sister of taxon1, taxon2, and taxon3, respectively.

During the training process, the four taxa in each quartet dataset were permutated to create 4! = 24 different orders, and each serves as an independent training sample, to ensure that the order of taxa in the dataset does not influence the phylogenetic inference. Two thousand trees randomly sampled from a total of 100,000 were used in each training epoch and were fed to the network in batches of 16 trees (each with 24 permutated samples). In the same batch, all 16 sequence alignments were resampled to contain the same number of sites. The resampling started by first randomly picking one of the 16 alignments and counting its number of sites *n*_aa_. Then, for each of the other 15 alignments, *n*_aa_ sites were sampled with replacement from the original sites of the alignment. Hence, the 16 alignments in the same batch were made to have equal sequence lengths to vectorize and accelerate the training process. We used the Adam optimizer with a learning rate of 0.001 and weight decay rate of 1×10^−5^. The cross-entropy loss function was used to measure the error of each prediction. Let us denote the three probabilities that a residual neural network outputs for the three possible topologies as *p*_1_, *p*_2_, and *p*_3_, respectively. The cross-entropy loss is calculated as 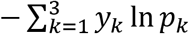, where *y_k_* is 1 if topology *k* is the truth and 0 otherwise. During validation and prediction, the order of the four taxa in each quartet dataset was also permutated to produce 24 alignments of the same sequences but with different orders of the four taxa. Each resultant alignment was subject to residual network prediction of three probability values, which provided the most likely topology for the alignment. The final prediction for the dataset was decided by the majority vote of the 24 predicted topologies of the 24 alignments made from the dataset.

### Tree and sequence simulation schemes

Each simulated tree was formulated as a PhyloTree object implemented by the Python package ete3 v3.1.1 (Huerta-Cepas, et al. 2016). To generate quartet datasets used in all training and validation processes and the datasets in **Table 1**, **Fig. 1**, and **Fig. 2**, we first generated a large tree with more than four taxa, and then pruned the tree to four taxa. Large tree topologies were randomly generated by the populate() method in ete3, while branch lengths were randomly assigned according to different distributions. When pruning a large tree into an LBA quartet tree, we first found an “LBA” tree for each internal node as follows. We identified the farthest leaf and the nearest leaf for each of the two children of this internal node; the four leaves form the four taxa of a quartet tree. A branch ratio statistic was calculated for the quartet tree, defined as (internal branch length + length of the longer of the two short branches) / length of the shorter of the two long branches. This process was repeated for all internal nodes, and the resultant quartet tree with the smallest branch ratio was retained as the pruned LBA tree. For trees in **Fig. 3** and those used in time benchmarking (testtime_01 in **Table S2**), the five branch lengths in a quartet tree were directly assigned without the process of pruning.

Sequences on a tree were simulated from more ancient to more recent nodes, starting at the root. On each branch of a tree, substitutions occurred at each individual site following a continuous-time Markov Chain in the fixed time represented by the branch length. When a substitution took place, the amino acid state was changed to another state with a probability proportional to the rate value defined in the transition rate matrix ***Q***. As stated in the main text, ***Q*** = ***S*Π**, where **Π** is the diagonal matrix of ***π*** (Yang 2006) and ***S*** is the exchangeability matrix. We modelled two aspects of heterogeneities: rate variation and profile difference. At each site, a relative evolutionary rate *r* was factored into the branch lengths of the tree for the site, and a site-specific equilibrium frequency profile ***π*** was assigned. Across all sites, *r* followed a gamma distribution. For simulating realistic ***π***, we summarized 4,986 site-specific equilibrium frequency profiles from 16 sequence alignments of nuclear and organelle proteins involving hundreds of taxa per alignment that were assembled in a previous study (Breen, et al. 2012). To decide the amino acid profile of a site in our simulation, one of the 4986 real profiles was picked randomly, which was then used as a base measurement vector parameter for a Dirichlet distribution. From this distribution with scale parameter 10, we then generated a random sample, which represents a new profile correlating with the real profile defining the Dirichlet distribution, but with variation. The Dirichlet distribution was chosen because it is a Bayesian conjugate of the multinomial distribution, which specifies the sampling probability of each amino acid at the focal site. The scale parameter was chosen to enable considerable variation of the profile, allowing more diverse evolution processes to be simulated. In addition to site heterogeneity, we added time heterogeneity to the simulation by changing the site-specific rate *r* and profile ***π*** of each site based on *r* and ***π*** of the root node, before simulating each branch. The time heterogeneity of rate variation (heterotachy) was realized by randomly selecting a fraction (*f*) of all sites and shuffling the *r* values among these sites. To create time heterogeneity of the amino acid profile at a site, we conducted the following operation on the profile vector of each amino acid site. First, two amino acid states were randomly selected. Second, their frequencies in the profile were swapped. Third, this process was repeated *n* times for each site. When heterogeneity existed in the simulation of a tree, there was a certain probability *p*_h_ for each branch to contain the time heterogeneity. If a branch was decided to be heterogeneous according to *p*_h_, both the heterotachy and frequency swap were performed.

Variable parameters in simulating a tree (before pruning to a quartet tree) include: number of taxa *M*, branch lengths *B*, number of amino acid sites *N*_aa_, exchangeability matrix ***S***, shape parameter *α* of the gamma distribution of the relative rate *r*, probability *p*_h_ with which a branch is heterogeneous, proportion of sites subject to rate shuffling *f*, and the number of profile swap operations *n* for each site. For each dataset simulated in this study, the values or distributions of these variable parameters are show in **Table S2**.

### Existing phylogenetic methods

NJ and MP inferences were conducted by MEGA -CC 10 (Kumar, et al. 2018). For NJ, the JTT model combined with a gamma distribution of rate variation (alpha parameter = 1) was used in calculating distances. For MP, tree searches was set as SPR. Ten initial trees and search level 1 were specified. ML inferences was conducted by RAxML v8.2.11 (Stamatakis 2014) and PhyML 3.1 v20120412 (Guindon, et al. 2010). For RAxML, the model was set to be “PROTCATAUTO”; for PhyML, the options used was “-d aa --sequential -n 1 -b 0 --model LG - f e --pinv e --free_rates --search BEST -o tl”. Bayesian inferences were performed by MrBayes pre-3.2.7 (Ronquist, et al. 2012). The LG transition matrix with a discrete gamma distribution of rate variation (with 4 rate categories) was used. MrBayes was used with two replicate runs, each with four chains running for 20,000 steps and a sampling frequency of every 50 steps. After 5,000 steps of the burn-in stage, the 50% majority-rule consensus tree was summarized for each inference as the output.

### Real phylogenetic datasets

The red fluorescent protein dataset was generated from an experimental phylogeny created by random mutagenesis PCR (Randall, et al. 2016). Amino acid sequences were extracted from Supplementary Note 1 of the study. From all sets of four taxa in the tree (**Fig. S2a**), 1,000 were randomly sampled to form quartet tree test datasets. All 1,000 datasets had an alignment length of 225 amino acids.

The 2,684 mammalian protein sequence alignments were downloaded from OrthoMaM v10 (Scornavacca, et al. 2019), by querying the database for genes present in all 24 species (**Fig. S2b**). For the first dataset, four species from the 24 species were randomly selected for each gene. If the ungapped alignment of this gene for the four taxa contained more than 50 amino acids, this alignment was retained, resulting in a total of 2,661 quartet trees. For the second dataset, all 2,684 alignments of 24 species were concatenated. Then, 2,000 quartet alignments were generated from this large alignment, by first randomly picking four species and then randomly sampling 100–3,000 (not necessarily consecutive) amino acid sites without replacement from all sites.

The seed plant dataset was downloaded from http://datacommons.cyverse.org/browse/iplant/home/shared/onekp_pilot/PNAS_alignments_trees/species_level/alignments/FAA/FAA.604genes.trimExtensively.unpartitioned.phylip (Wickett, et al. 2014). We then compiled 1,000 quartet alignments by first randomly picking four species that do not form polytomy (**Fig. S2c**) and then randomly sampling 100–3,000 (not necessarily consecutive) sites without replacement from all sites.

## ACKNOWLEDGEMENTS

This project was supported by the Michigan Institute for Computational Discovery & Engineering Catalyst Grant to J.Z. and Y.G. and U.S. National Institutes of Health grant GM103232 to J.Z. We thank the support of NVIDIA Corporation with the donation of the Titan Xp GPU used in this research.

## Supplementary materials

Supplementary materials include:

Tables S1–;S2

Supplementary figure legends

Figures S1–;S2

**Table S1.**
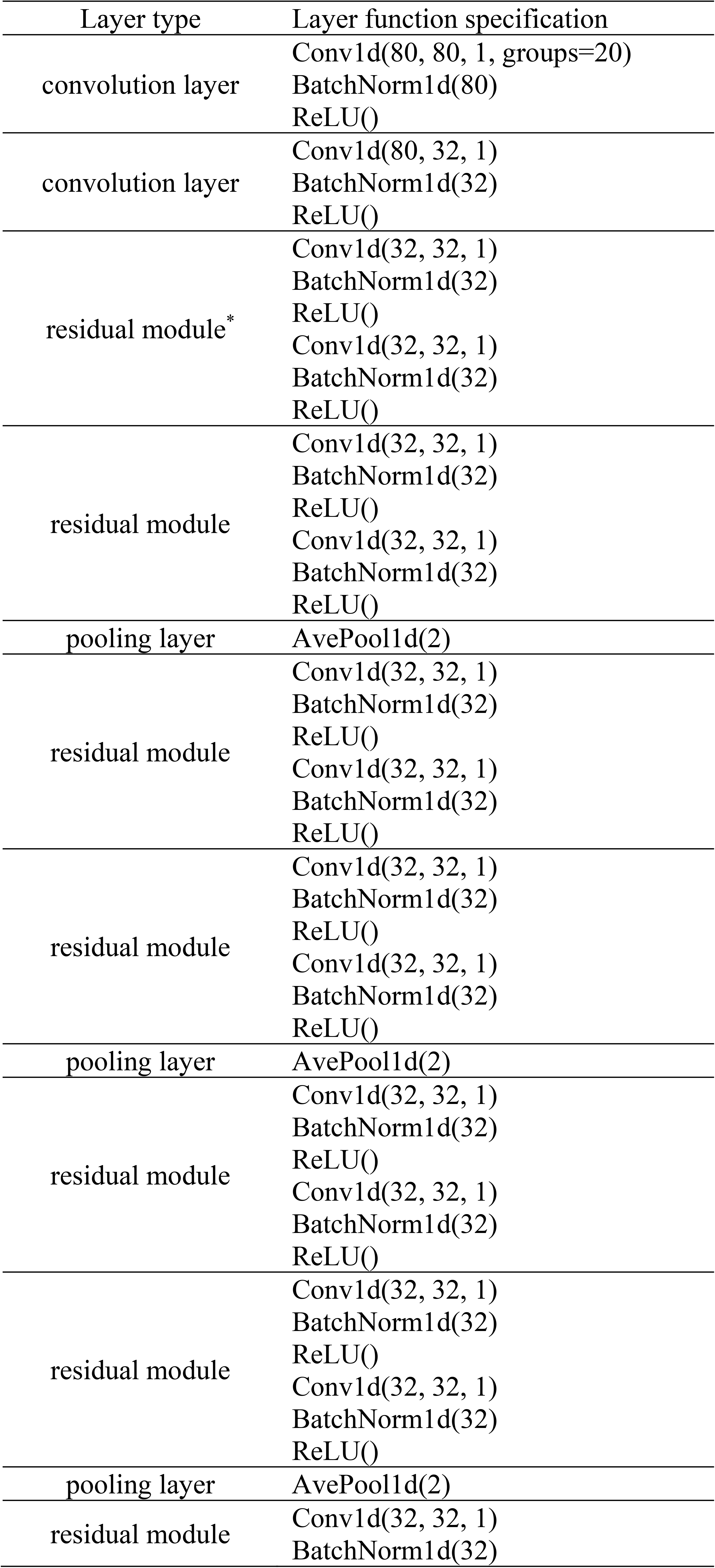
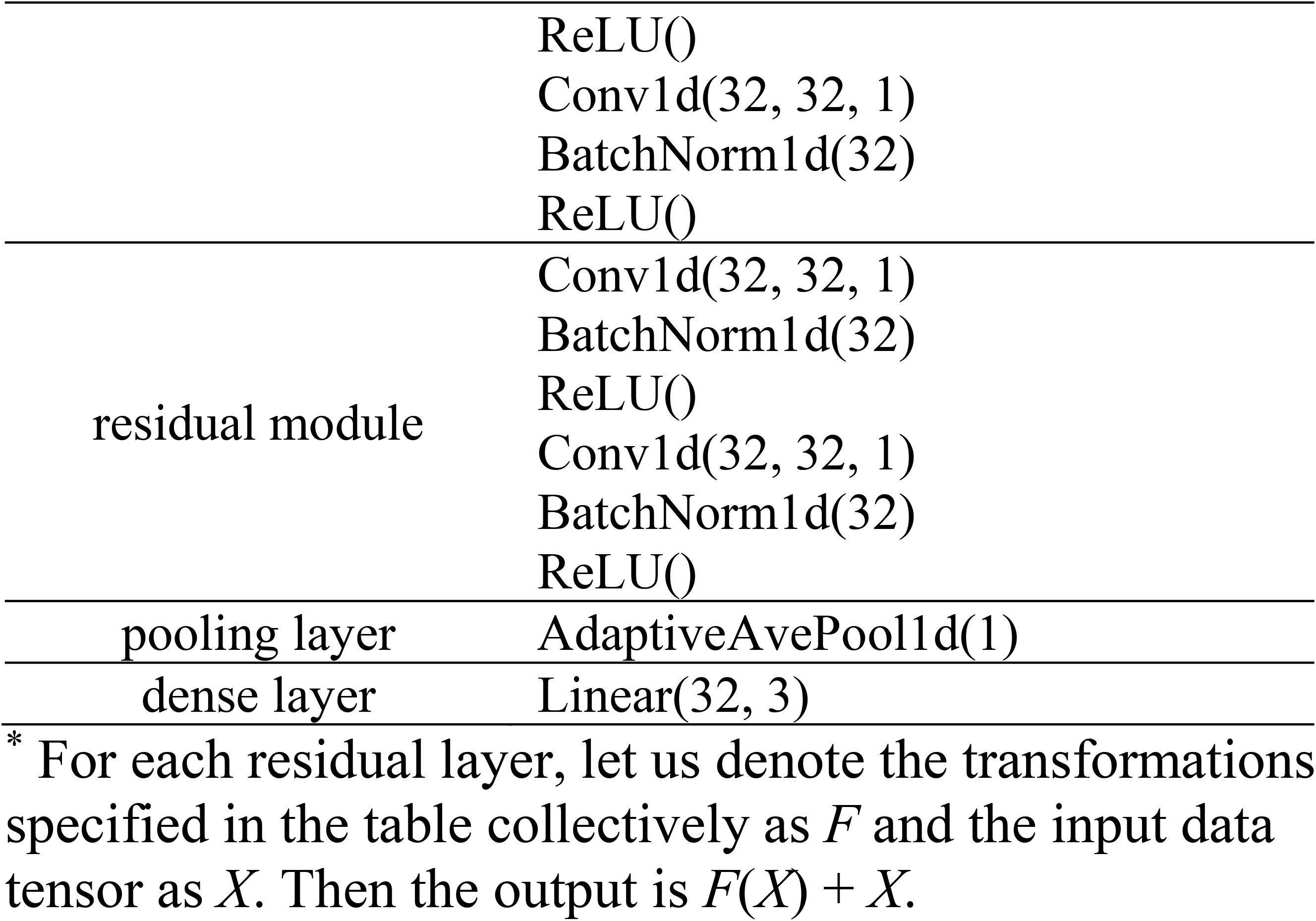
Residue neural network structure in PyTorch layer functions.

**Table S2.** Parameter settings for difference datasets / simulation schemes. See Materials and Methods for detailed explanation of different parameters.

| Dataset | $M$ | $B$ | $B$ | $N_{aa}$ | $S$ | $\alpha$ | $p_h$ | $f$ | $n$ |
| --- | --- | --- | --- | --- | --- | --- | --- | --- | --- |
|  | number of taxa | internal branches | external branches | site number | exchangeability<sup>a</sup> | gamma shape | heterogeneity | swap fraction | swap number |
| training1 (testing1_mixed) | Uniform(5, 105)<sup>b</sup> | Uniform(0.02, 1.02) | Uniform(0.02, 1.02) | Uniform(100, 3000) | random | Uniform(0.05, 1) | 0.9 | Uniform(0, 1) | Uniform(10, 20)<sup>b</sup> |
| testing1_nolba | Uniform(5, 105) | Uniform(0.02, 1.02) | Uniform(0.02, 1.02) | Uniform(100, 3000) | random | Uniform(0.05, 1) | 0.9 | Uniform(0, 1) | Uniform(10, 20) |
| testing1_lba | Uniform(5, 105) | Uniform(0.02, 1.02) | Uniform(0.02, 1.02) | Uniform(100, 3000) | random | Uniform(0.05, 1) | 0.9 | Uniform(0, 1) | Uniform(10, 20) |
| training2 (testing2_mixed) | Uniform(5, 105) | Gamma(1, 0.1) | Gamma(2, 0.1) | Uniform(100, 3000) | random | Uniform(0.05, 1) | 0.9 | Uniform(0, 1) | Uniform(0, 20) |
| testing2_nolba | Uniform(5, 105) | Gamma(1, 0.1) | Gamma(2, 0.1) | Uniform(100, 3000) | random | Uniform(0.05, 1) | 0.9 | Uniform(0, 1) | Uniform(0, 20) |
| testing2_lba | Uniform(5, 105) | Gamma(1, 0.1) | Gamma(2, 0.1) | Uniform(100, 3000) | random | Uniform(0.05, 1) | 0.9 | Uniform(0, 1) | Uniform(0, 20) |
| training3 (testing3_mixed) | Uniform(5, 105) | Uniform(0.02, 1.02) | Uniform(0.02, 1.02) | Uniform(100, 3000) | random | Uniform(0.05, 1) | Uniform(0, 1) | Uniform(0, 1) | Uniform(0, 20) |
| testing3_nolba | Uniform(5, 105) | Uniform(0.02, 1.02) | Uniform(0.02, 1.02) | Uniform(100, 3000) | random | Uniform(0.05, 1) | Uniform(0, 1) | Uniform(0, 1) | Uniform(0, 20) |
| testing3_lba | Uniform(5, 105) | Uniform(0.02, 1.02) | Uniform(0.02, 1.02) | Uniform(100, 3000) | random | Uniform(0.05, 1) | Uniform(0, 1) | Uniform(0, 1) | Uniform(0, 20) |
| test_branch_00 | 20 | Uniform(0.02, 0.2) | Uniform(0.02, 0.2) | 1000 | random | Uniform(0.05, 1) | 0.9 | Uniform(0, 1) | Uniform(0, 20) |
| test_branch_01 | 20 | Uniform(0.2, 0.4) | Uniform(0.2, 0.4) | 1000 | random | Uniform(0.05, 1) | 0.9 | Uniform(0, 1) | Uniform(0, 20) |
| test_branch_02 | 20 | Uniform(0.4, 0.6) | Uniform(0.4, 0.6) | 1000 | random | Uniform(0.05, 1) | 0.9 | Uniform(0, 1) | Uniform(0, 20) |
| test_branch_03 | 20 | Uniform(0.6, 0.8) | Uniform(0.6, 0.8) | 1000 | random | Uniform(0.05, 1) | 0.9 | Uniform(0, 1) | Uniform(0, 20) |
| test_branch_04 | 20 | Uniform(0.8, 1.0) | Uniform(0.8, 1.0) | 1000 | random | Uniform(0.05, 1) | 0.9 | Uniform(0, 1) | Uniform(0, 20) |
| test_branch_05 | 20 | Uniform(1.0, 2.0) | Uniform(1.0, 2.0) | 1000 | random | Uniform(0.05, 1) | 0.9 | Uniform(0, 1) | Uniform(0, 20) |
| test_seqlen_00 | 20 | Uniform(0.02, 1.02) | Uniform(0.02, 1.02) | Uniform(100, 200) | random | Uniform(0.05, 1) | 0.9 | Uniform(0, 1) | Uniform(0, 20) |
| test_seqlen_01 | 20 | Uniform(0.02, 1.02) | Uniform(0.02, 1.02) | Uniform(200, 500) | random | Uniform(0.05, 1) | 0.9 | Uniform(0, 1) | Uniform(0, 20) |
| test_seqlen_02 | 20 | Uniform(0.02, 1.02) | Uniform(0.02, 1.02) | Uniform(500, 1000) | random | Uniform(0.05, 1) | 0.9 | Uniform(0, 1) | Uniform(0, 20) |
| test_seqlen_03 | 20 | Uniform(0.02, 1.02) | Uniform(0.02, 1.02) | Uniform(1000, 3000) | random | Uniform(0.05, 1) | 0.9 | Uniform(0, 1) | Uniform(0, 20) |
| test_seqlen_04 | 20 | Uniform(0.02, 1.02) | Uniform(0.02, 1.02) | Uniform(3000, 10000) | random | Uniform(0.05, 1) | 0.9 | Uniform(0, 1) | Uniform(0, 20) |
| test_heterotachy_00 | 20 | Uniform(0.02, 1.02) | Uniform(0.02, 1.02) | 1000 | random | Uniform(0.05, 1) | 1.0 | 0.0 | 0 |
| test_heterotachy_01 | 20 | Uniform(0.02, 1.02) | Uniform(0.02, 1.02) | 1000 | random | Uniform(0.05, 1) | 1.0 | 0.25 | 0 |
| test_heterotachy_02 | 20 | Uniform(0.02, 1.02) | Uniform(0.02, 1.02) | 1000 | random | Uniform(0.05, 1) | 1.0 | 0.5 | 0 |
| test_heterotachy_03 | 20 | Uniform(0.02, 1.02) | Uniform(0.02, 1.02) | 1000 | random | Uniform(0.05, 1) | 1.0 | 0.75 | 0 |
| test_heterotachy_04 | 20 | Uniform(0.02, 1.02) | Uniform(0.02, 1.02) | 1000 | random | Uniform(0.05, 1) | 1.0 | 1.0 | 0 |
| testlba_F_h0 | 4 | varies<sup>c</sup> | varies | 2000 | wag.dat | 1 | 0.0 | N.A. | N.A. |
| testlba_F_h1 | 4 | varies | varies | 2000 | wag.dat | 1 | 1.0 | 1.0 | 0 |
| testtime_01 | 4 | 1.0 | 1.0 | 2000 | random | 0.5 | 0.0 | N.A. | N.A. |

<sup>a</sup> When $S$ is “random”, each simulated tree in the dataset is assigned an exchangeability matrix $S$ randomly chosen from {“cpREV64.dat”, “dayhoff.dat”, “jones.dat”, “lg.dat”, “wag.dat”, “mtArt.dat”, “mtmam.dat”, “mtREV24.dat”, “MtZoa.dat”} from the data files implemented in PAML 4.9.
<sup>b</sup> Uniform distributions for $M$ and $n$ only take integer values with inclusive lower bounds and exclusive upper bounds.
<sup>c</sup> Different datasets in “testlba” series of datasets have different branch length assignments.

## SUPPLEMENTARY FIGURE LEGENDS

**Fig. S1.**
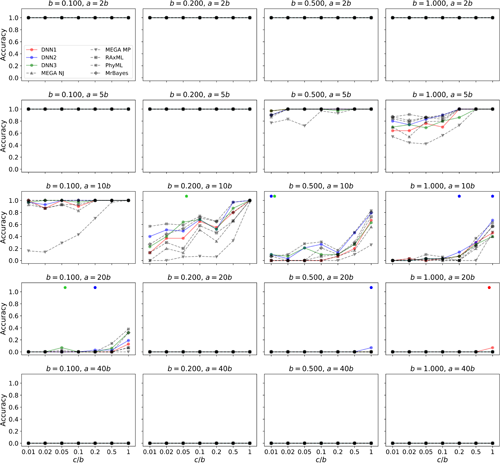
Performances of residual network predictors and existing methods on LBA trees without heterotachy. Shown are proportions of 100 quartet trees correctly inferred by our predictors (shown by different colors) and the existing methods (shown by different grey symbols) under various combinations of the parameters *b, a*/*b*, and *c*/*b*. For each *c*/*b* level indicated on the *X*-axis, if a residual network predictor performs better than all existing methods, a pentagon of the corresponding color is drawn on the top of the panel.

**Fig. S2.**
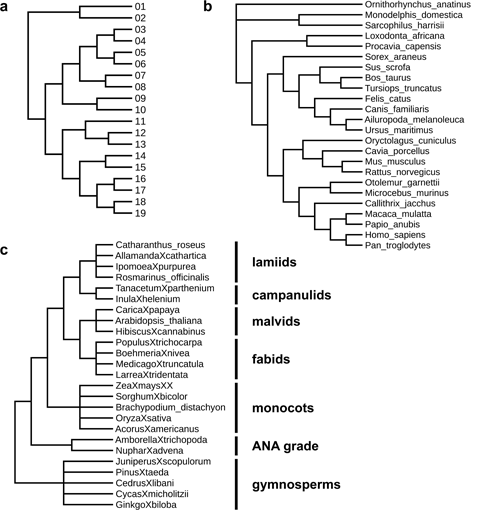
True trees assumed for three actual phylogenetic datasets. (**a**) The actual cladogram of the lab evolved red fluorescent proteins (Randall, et al. 2016). (**b**) The presumed cladogram of 24 mammalian species. (**c**) The presumed cladogram of 25 seed plant species. Black vertical lines indicate seven clades whose phylogenetic relationships are specified and used in comparing performances of different methods.

